# Expansion of functional human salivary acinar cell spheroids with reversible thermo-ionically crosslinked 3D hydrogels

**DOI:** 10.1101/2024.09.09.612117

**Authors:** Jose G. Munguia-Lopez, Sangeeth Pillai, Yuli Zhang, Amatzia Gantz, Dimitria B. Camasao, Showan N. Nazhat, Joseph M. Kinsella, Simon D. Tran

## Abstract

Xerostomia (dry mouth) is frequently experienced by patients treated with radiotherapy for head and neck cancers or with Sjögren’s syndrome, with no permanent cure existing for this debilitating condition. To this end, *in vitro* platforms are needed to test therapies directed at salivary (fluid-secreting) cells. However, since these are highly differentiated secretory cells, the maintenance of their differentiated state while expanding in numbers is challenging. In this study, the efficiency of three reversible thermo-ionically crosslinked gels: 1) alginate-gelatin (AG), 2) collagen-containing AG (AGC), and 3) hyaluronic acid-containing AG (AGHA), to recapitulate a native-like environment for human salivary gland (SG) cell expansion and 3D spheroid formation was compared. Although all gels were of mechanical properties comparable to human SG tissue (∼11 kPa) and promoted the formation of 3D spheroids, AGHA gels produced larger (>100 cells/spheroid), viable (>93%), proliferative, and well-organized 3D SG spheroids while spatially and temporally maintaining the high expression of key SG proteins (aquaporin-5, NKCC1, ZO-1, α-amylase) for 14 days in culture. Moreover, the spheroids responded to agonist-induced stimulation by increasing α-amylase secretory granules. Here, we propose alternative low-cost, reproducible, and reversible AG-based 3D hydrogels that allow the facile and rapid retrieval of intact, highly viable 3D-SG spheroids for downstream applications.

## Introduction

Salivary glands (SGs) are exocrine organs responsible for the production and secretion of saliva, a complex, multifunctional, extracellular fluid that maintains oral health.^1,2^ Irreversible SG damage leads to hyposalivation, manifesting itself as dry mouth symptoms (xerostomia) in patients treated with radiotherapy for head and neck cancers and in patients with an autoimmune disease, such as Sjögren’s syndrome.^1–3^ Irreversible xerostomia is mainly caused by a loss of acinar cell number and their functionality; acinar cells are the principal site in saliva secretion.^2,4^ Currently, there is no permanent treatment available for patients with damaged salivary epithelial (acinar) cells or those replaced by fibrotic tissues. Existing pharmacological options largely involve saliva-stimulating drugs, relying on the presence of surviving acinar cell populations. Palliative care represents an alternative strategy for xerostomia, providing temporary relief from symptoms, but lacks long-term therapeutic benefits.^1,2^

The bioengineering of fully functional SG models is particularly demanding due to complex tissue physiology and challenges in mimicking the highly coordinated actions of multiple cell types (acinar, ductal, and myoepithelial cells) *in vitro.*^1,2^ Furthermore, it is imperative to establish a carefully orchestrated microenvironment that facilitates the attachment, proliferation, and reorganization of these specialized cells to closely mimic the native tissue. Among the various critical factors, soluble cues, extracellular matrix (ECM) analogs, and mechanical properties stand out as critical parameters for the successful construction of an artificial SG. Despite significant efforts from the scientific community to devise new tools for the expansion and creation of SG organoids, the refinement of culture conditions and three-dimensional (3D) platforms, particularly using biomimetic hydrogels, remains an ongoing area of development. A substantial body of literature on SG bioengineering continues to advocate for the use of animal-derived matrices in cell culture, which may potentially restrict the translational potential of these outcomes in the long term.

Hydrogels have been extensively used as platforms for biomedical applications due to their ability to bioengineer an artificial 3D environment that mimics the native mammalian niche.^1,5^ Among the naturally derived hydrogels, alginate and gelatin have been widely utilized as ECM analogs in 3D *in vitro* platforms, either as standalone scaffolding materials^6,7^ or in the form of composite gels for organoid/spheroid formation.^8–10^ In terms of their attributes in composite hydrogels, alginate enables mechanical reinforcement via ionic crosslinking, thus controlling gel stiffness, whereas gelatin provides rapid thermal gelation while supplying potential adhesive sites for cell binding, thereby forming focal adhesion complexes and reinforcing essential cell-ECM interactions for SG cell reorganization in 3D.^10–12^ We have demonstrated the potential application of alginate-gelatin (AG) composite hydrogels as printable 3D *in vitro* platforms for cancer spheroid formation.^8–10,13^ Collagen, which is the most abundant ECM protein found in mammalians and one of the major constituents of the basement membrane,^2,14^ provides cell adhesion sites (*i.e.,* arginine-glycine-aspartic acid that enable cell-ECM interactions leading to cell adhesion), and is involved in SG acinar cell differentiation during organogenesis^15^, thereby playing a critical role in SG branching morphogenesis^1^. Collagen is mainly used in combination with other biomaterials when investigated for SG bioengineering, especially based on its presence in animal-derived matrix mimetics,^16^ thus forming a non-reversible, stable matrices/substrates.

Hyaluronic acid (HA) is another biopolymer that has been explored for SG tissue engineering. HA is a non-sulfated glycosaminoglycan ECM component, expressed during SG organogenesis, which has been utilized to facilitate the formation of SG spheroids and the growth of SG epithelial cells.^17–19^ This is attributed to its pivotal role in cell adhesion, migration, and morphogenesis by binding with cell surface receptors, such as CD44 (expressed by SG acinar cells^4,20^) and RHAMM.^21,22^ Nevertheless, although the utilization of chemically modified (*e.g.,* thiolate, acrylate, bioactive basement membrane-derived peptides),covalently crosslinked hydrogels has been documented in SG tissue engineering,^2,18,23^ the potential of HA as a freely suspended moiety in combination with other naturally derived composite hydrogels remains unexplored.

Because the matrix composition plays an important role in spheroid formation, in this study, we evaluated the potential of three hydrogels of varying ECM-mimicking components in enhancing the proliferation and formation of 3D human SG acinar spheroids. Specifically, we compared the potential of AG hydrogels to those incorporated with either collagen (AGC) or HA (AGHA) for SG acinar cell reorganization into 3D spheroids, *in vitro*. Collagen and HA were added to evaluate their impact on SG acinar cell attachment, proliferation, viability, and self-assembly reorganization into 3D spheroids by either increasing the cell adhesion sites (proportionated by AGC) or by promoting the CD44 receptor-mediated acinar cell reorganization (using AGHA), respectively. Fig. 1 shows the workflow of the hanging drop approach used for 3D spheroid production and evaluation in these hydrogels. The mechanical properties of the hydrogels were initially characterized, followed by an assessment of their performance in supporting human SG cell proliferation, viability, and spheroid formation for up to 14 days in culture. Having identified that HA incorporation promoted the formation of highly viable, proliferative, and large acinar spheroids, AGHA hydrogels were selected as the 3D platform to evaluate the expression and localization of key SG proteins and spheroid functionality, as well as fostering human-derived salivary functional units (SFU). It is anticipated that the reversible nature of AG-based hydrogels will allow for easy retrieval of highly viable SG spheroids and organoids by a simple ion chelation process at 37°C for downstream applications, without compromising the integrity of the biological entities.

**Fig. 1.**
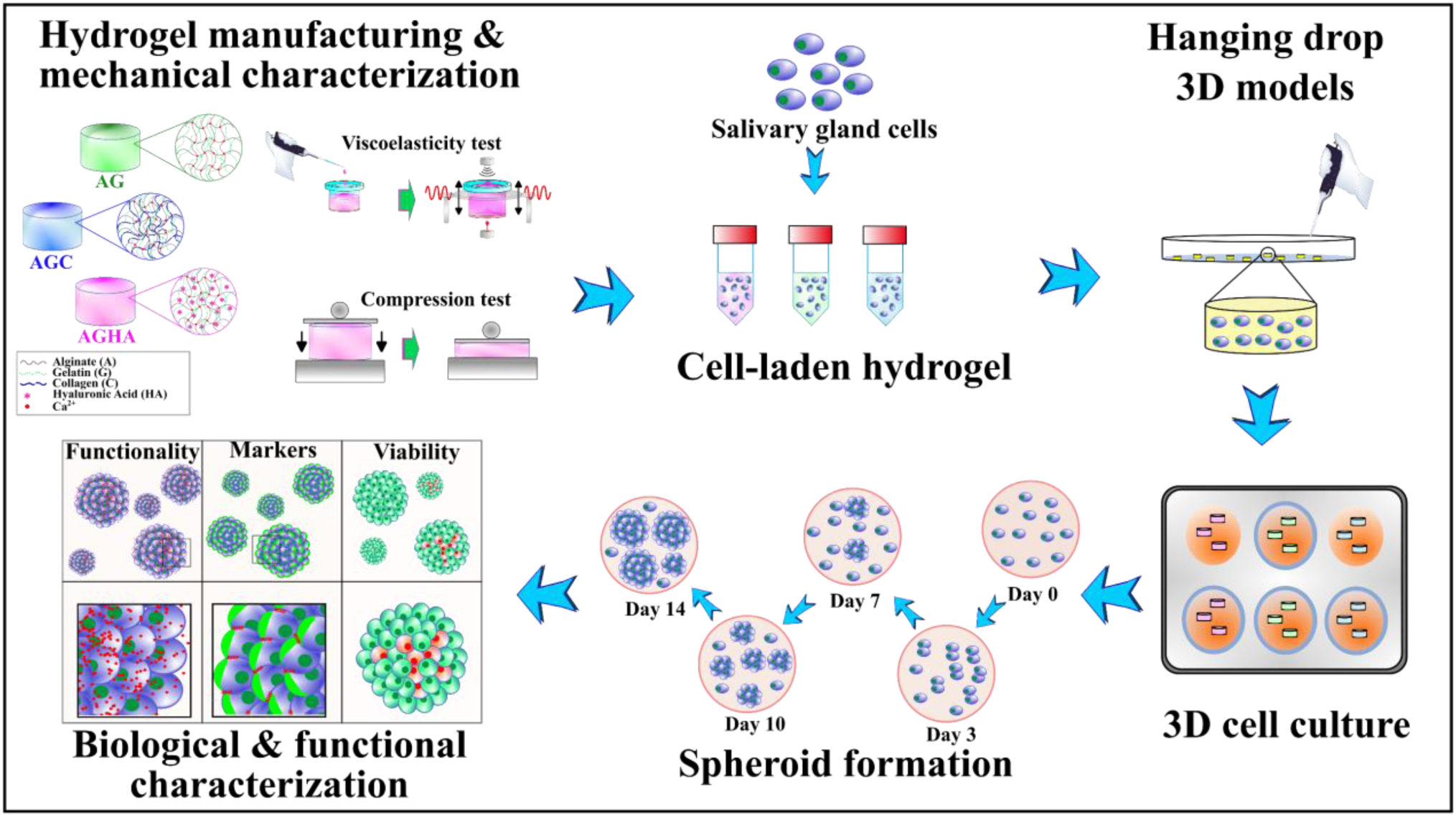
Schematic depiction of the SG 3D culture workflow as spheroid forming platforms using the hanging drop technique. AG, AGC, and AGHA hydrogels were synthesized and mechanically characterized. Next, human SG cells (cultured in the 2D platform) were harvested with trypsin, counted, and mixed separately within hydrogels. Then, drops of 30 µl (3D models) were created by the hanging drop method. The 3D models were thermogelled, ionically crosslinked, and cultured for 14 days. The formation of spheroids was tracked through time by confocal microscopy imaging. Based on hydrogel performance in promoting the formation of highly viable, proliferative, and large acinar spheroids, the AGHA gel was selected to evaluate the expression and localization of key SG proteins and functionality in the produced spheroids. AGHA gels were used to expand and foster functional SFU.

## Results

### Biologically derived hydrogels show mechanical properties comparable to human SG tissues

The mechanical properties of the hydrogels were analyzed by measuring the changes in viscoelastic properties through gelation, as well as by carrying out micro-compression testing on as-formed gels. A contactless, non-destructive technique^24^ measured the temporal change in shear storage modulus (G’, Fig. 2 a) in real-time, as governed by the thermogelation of gelatin. Gelation occurred at a similar rate for all three formulations, as indicated by the increase in G’ during the first 10 min and until reaching their maximum G’ (plateau). While HA-containing hydrogels were of lower G’ values than those of AG and AGC (Fig. 2 b), there were no statistically significant (p>0.05) differences (Fig. 2 c) in maximum average G’ values, which ranged from ∼195 to 220 Pa. On the other hand, ionic crosslinking of the alginate component resulted in an increase in G’ values and significant (p<0.05) differences among the three hydrogel compositions (Fig. 2 c), where AG showed the highest maximum average G’ value (1.82±0.09 kPa), followed by AGC (1.74±0.07 kPa), and AGHA (1.57±0.14 kPa). By assuming that the hydrogels are isotropic, homogeneous and incompressable, with a Poisson’s ratio (ν) of 0.5,^10,25,26^ storage elastic modulus (E’) values were approximated through the equation: *E* = 2(1 + *v*)*G*. ^27^ Calculated E’ values for the thermogelled gels were 0.64±0.07 kPa, 0.63±0.08 kPa, and 0.59±0.05 kPa for AG, AGC, and AGHA, respectively, whereas those for the ionically crosslinked gels, were 5.43±0.28 kPa, 5.22±0.22 kPa, and 4.72±0.44 kPa for AG, AGC and AGHA, respectively.

**Fig. 2.**
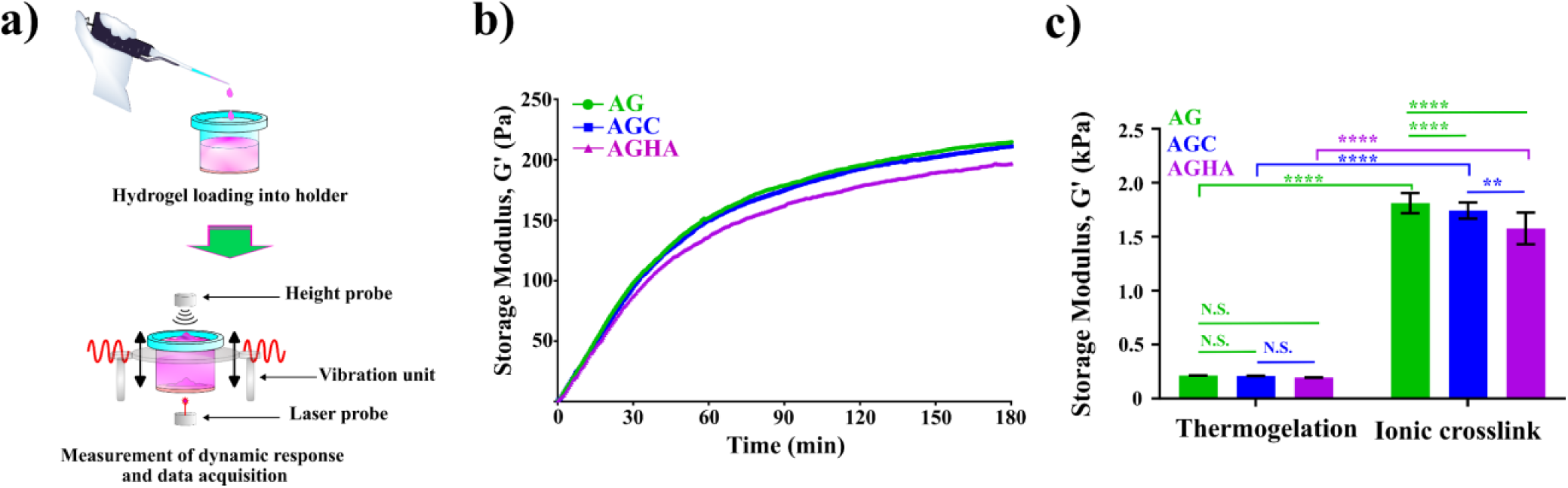
Viscoelastic properties of the thermo-ionically crosslinked hydrogels. a) Schematic images of the ElastoSens Bio2 instrument used to characterize gelation through measurment of viscoelastic properties by a contactless, non-destructive approach. b) Evolution of shear storage modulus (G’) as a function of time, and c) maximum average G’ of hydrogels based on gelatin gelation curves (extracted from b) and ionically crosslinked gels. Data presented as mean ± SD, n≥3, N.S. not significant, **p<0.01, ****p<0.0001.

Micro-compression testing was used to compare the compressive modulus at low strain (0.05-0.10) of as-formed gels (Fig. 3 a), with representative stress-strain curves demonstrating typical densification trends (Fig. 3 b). The thermogelled hydrogels did not show significant (p>0.05) differences in compressive modulus with values of ∼1.5 kPa (Fig. 3 c). Ionic crosslinking of the alginate component generated gels of higher moduli (Fig. 3 d-f), with AG, AGC, AGHA demonstrating values of 10.81±1.42 kPa, 11.88±1.45 kPa, 10.64±2.33 kPa, respectively, and no significant (p>0.05) differences between the gels (Fig. 3 f).

**Fig. 3.**
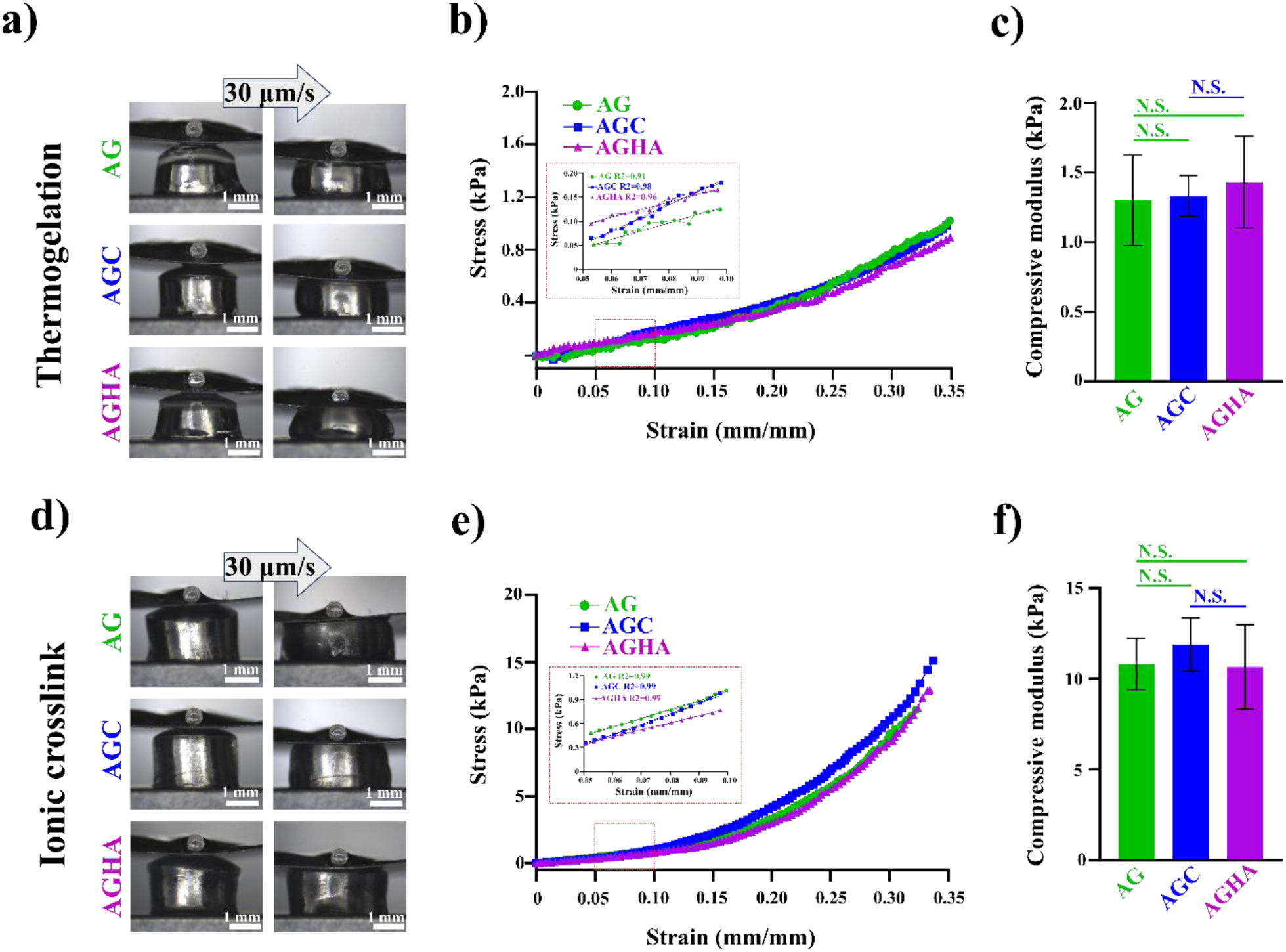
Micro-compression testing of as-formed hydrogels. a) & d) Representative images of micro-compression testing of thermo-gelled and ionically crosslinked hydrogels at a displacement rate of 30 µm/s. b) & e) Typical compressive stress-strain curves of thermo-gelled and crosslinked hydrogels, respectively; inserts show the linear region at low strains (0.05-0.10). c) & f) Compressive modulus values calculated from the stress-strain slope at low strains (red dotted boxes inserts in b & e). Data presented as mean ± SD, n≥3, N.S. not significant, **p<0.01, ****p<0.0001.

### HA-containing hydrogels promote the self-assembly of highly viable, proliferative, and sizable 3D NS-SV-AC spheroids

Confocal microscopy imaging was used to assess the viability, metabolic activity, as well as spheroid formation and growth of the human SG acinar cell line NS-SV-AC, as a function of time in culture within the gels (Fig. 4). Spheroid formation initiated within 3 days in culture in all hydrogels, which contoured into larger, well-compacted, and 3D spherical structures containing more than 50 cells/spheroid by day 7, that further increased to >100 cells/spheroid at day 10 in culture (Fig. 4 a-c). Live/dead staining revealed that spheroids in AG and AGC contained a large number of cells stained as red (ethidium homodimer I), indicating cell death (Fig. 4 a & b, and Supplementary Fig. 1 a & b). In contrast, no cell death was found in the core of sizeable 3D structures growing in AGHA (Fig. 4 c, Supplementary Fig. 1 c). The presence of dead cells in the core of spheroids was higher in AGC, followed by AG and AGHA (Fig. 4 a-c). The viability of spheroids in all gels increased during the first 3 days of culture, reaching a plateau by day 10. At day 14, the viability of spheroids was significantly (p<0.05) higher in AGHA compared to the other gels (Fig. 4 d, Table 1).

**Fig. 4.**
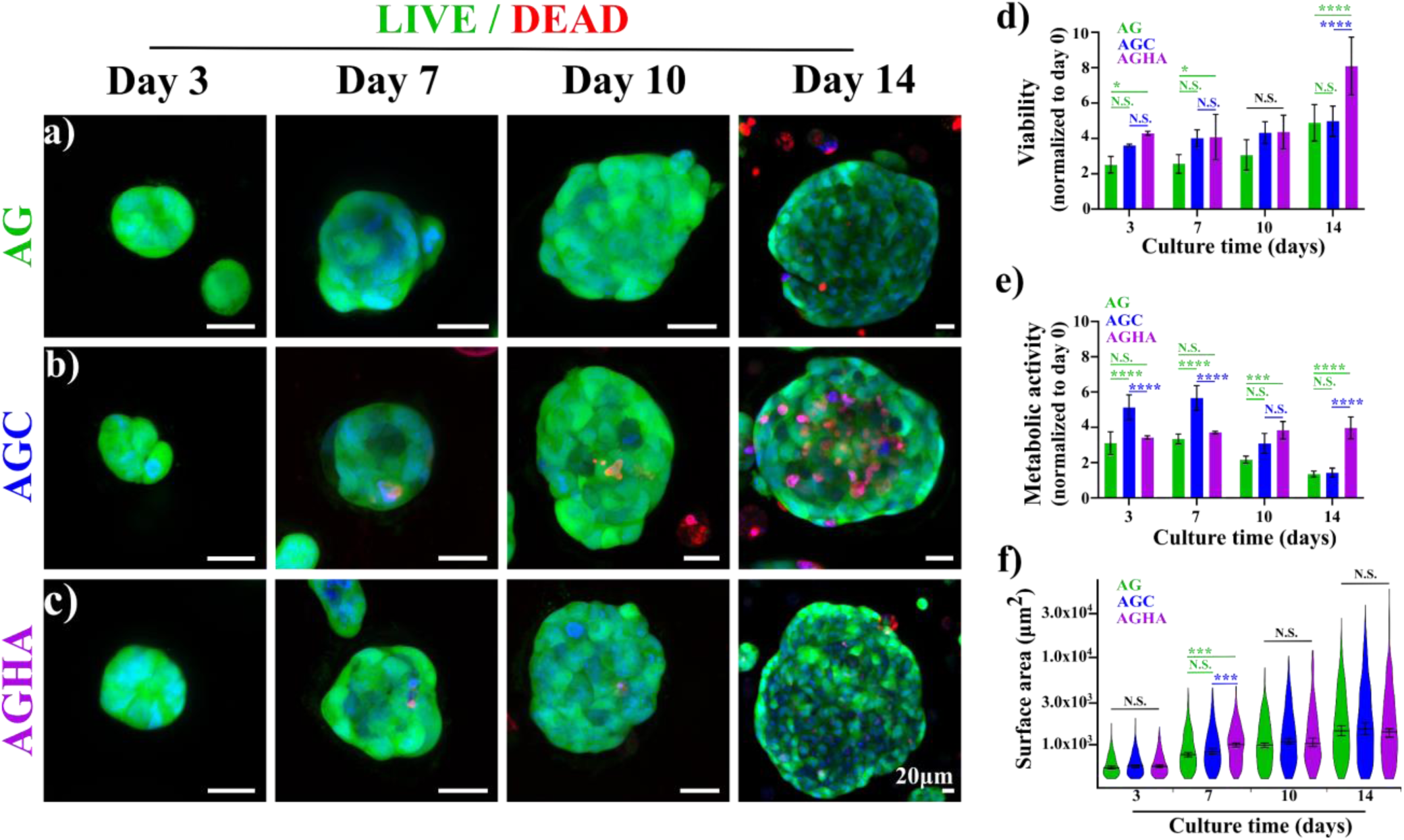
3D NS-SV-AC spheroid formation in hydrogels. Cells were seeded into a) AG, b) AGC, or c) AGHA hydrogels and cultured for 14 days. Live/dead assay confocal imaging was performed to determine spheroid morphology (a-c), viability (d), and size (f). Metabolic activity was measured using WST-1 assay (e). Green, represents live cells; red, dead cells; blue, nuclear staining. Magnification/scale bar: x20/20 μm. Data presented as mean ± SD, n≥3, N.S., no significant, *p<0.05, **p<0.01, ****p<0.0001.

**Table 1.** Summary of hydrogel performance as a 3D cell expansion platform.

|  |  | Hydrogel composition |  |  |
| --- | --- | --- | --- | --- |
|  |  | AG | AGC | AGHA |
| **Viability (%)** | *Day 3* | 84.20±3.74 | 90.87±0.76 | 90.57±0.65 |
|  | *Day 7* | 85.75±2.89 | 84.20±1.75 | 89.71±1.71 |
|  | *Day 10* | 84.26±2.42 | 87.78±0.41 | 90.41±1.24 |
|  | *Day 14* | 87.82±2.56 | 81.00±1.46 | 93.21±1.96 |
| **Maximum spheroid surface area (µm<sup>2</sup>)** | *Day 3* | 1,742.70 | 1,996.08 | 1,582.03 |
|  | *Day 7* | 4,449.46 | 4,461.82 | 4,653.40 |
|  | *Day 10* | 7,601.17 | 10,351.19 | 12,050.64 |
|  | *Day 14* | 28,056.36 | 41,225.55 | 61,204.89 |
| **Dead cells in spheroid core (day 14)** |  | Few cells | Yes | No |
| **<sup>a)</sup>Performance** |  | 3 | 3 | 5 |

The viability percentage was determined by culture per day per condition and not normalized with day 0 values. ND: not determined.
<sup>a)</sup>Hydrogel and culture performance as cell expansion system: arbitrary scale considering viability, metabolic activity (proliferation), spheroid formation, spheroid size, and presence of dead cells in their core. 1: very bad, 2: bad, 3: regular, 4: good, 5: excellent.

The metabolic activity of the spheroids was measured as an indicator of their proliferation, which demonstrated an increase of almost 4-fold in AGHA and AG hydrogels during the first 3 days of culture. This level of metabolic activity of the spheroids growing in AGHA was maintained between days 7 and 14, whereas the activity of those in AG gels significantly (p<0.05) decreased after day 10 in culture. In contrast, while spheroid metabolic activity in AGC gels was higher compared to the other gels during the first 7 days of culture - demonstrating up to ∼5-6 fold increase – this decreased rapidly thereafter. Both AG and AGC gels presented similar decreasing trends in activity after day 10 in culture (Fig. 4 e).

Based on its surface area, spheroid size may be classified as either small (400-10,000 µm^2^), medium (10,000-20,000 µm^2^), or large (>20,000 µm^2^).^8,9^ For analysis of spheroid size in this study, only live spheroids (green, calcein-AM dye) were considered. There was an increase in NS-SV-AC spheroid size within all gels as a function of time, where a range of small, medium, and large spheroids was found. Medium-sized spheroids contained >100 cells/spheroid,, and the most prominent structures found in gels ranged from 23,000 to 61,000 µm^2^, where AGHA promoted the formation of larger 3D structures followed by AGC and AG, respectively (Fig. 4 f, Table 1). Nonetheless, there were no statistically significant (p>0.05) differences in spheroid size between the gels after 10 days in culture (Fig. 4 f, Table 1).

Spheroid formation was further evaluated by cytoskeleton protein staining (F-actin) where confocal imaging revealed that all 3D structures within the gels were well-organized, spherically compact entities (Supplementary Fig. 2). Given together, the AGHA gels showed superior outcomes in terms of NS-SV-AC cell viability, spheroid formation and proliferation, and their reorganization into larger viable structures over the 14-day culturing period and where selected for further analysis in this study.

### NS-SV-AC spheroids show a robust expression of acinar cell-specific markers

RT-qPCR was used to compare the extent of expression of four essential salivary gland genes responsible for the secretion of saliva and salivary proteins: AQP5, tight junction protein zona occludens-1 (ZO-1), sodium-potassium-chloride cotransporter type 1 (NKCC-1), and salivary enzyme amylase (AMY-1) in NS-SV-AC cells growing in 2D, and 3D, when seeded into AGHA gels. AQP5, ZO-1, NKCC-1, and α-amylase showed significantly (p<0.05) higher expression in 3D compared to 2D (Fig. 5 a-d), suggesting that the 3D models promoted the expression of these acinar markers.

**Fig. 5.**
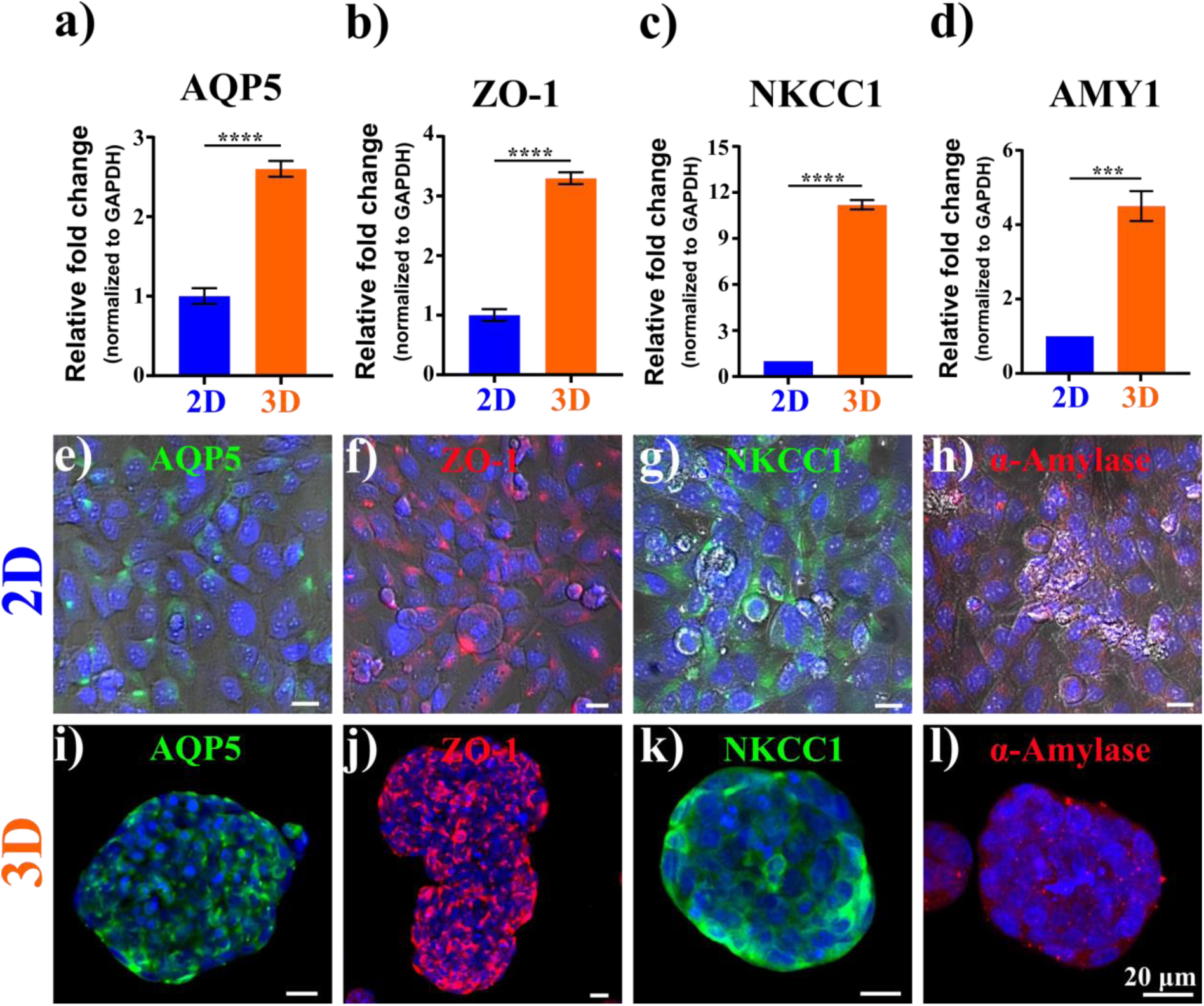
Analysis of key acinar cell markers in 3D NS-SV-AC spheroids. mRNA transcript levels were analyzed by RT-qPCR for acinar-specific markers expressed in 2D (plastic, 4 days in culture) and 3D platform (AGHA, 14 days in culture) (a-d). Data were normalized to GAPDH values. Data presented as mean ± SD, n≥3, ***p<0.001, ****p<0.0001. Immunodetection of acinar markers in NS-SV-AC cells growing in 2D (on plastic) and 3D platforms (in AGHA gels). 2D maximum projection of markers expressed by cells in 2D (e-h) or 3D (i-l) systems. For 2D culture images (e-h), the bright field was included in the merged images to increase the contrast of individual cells. Green, AQP5 or NKCC-1; red, ZO-1 or α-amylase; blue, nuclear staining. Magnification x20, scale bar 20 µm.

Immunodetection and localization of these acinar cell markers, along with PanCK, Col-IV, Pan-Laminin, Vimentin, and F-actin, showed that the 2D cultured NS-SV-AC cells expressed AQP5, ZO-1, NKCC-1, and α-amylase proteins at low levels (Fig. 5 e-h), with AQP-5 proteins localized only to the cellular cytoplasms (Fig. 5 e). The majority of the cells showed the tight junction-associated protein (ZO-1), but without apical-lateral localization (Fig. 5 f). Moreover, the distribution of NKCC-1 appeared to be similar in both cytoplasm and membrane (Fig. 5 g) and the presence of α-amylase was detected as small cytoplasmic granules (Fig. 5 h). PanCK was detected as a cytoskeleton protein, confirming the epithelial nature of NS-SV-AC cells (Supplementary Fig. 3 a). The extracellular proteins of Col-IV and Pan-Laminin were detected on the membrane/cell junction area (Supplementary Fig. 3 b & c, respectively), whereas Col-IV was present only in part of the NS-SV-AC population (Supplementary Fig. 3 b). Vimentin and F-actin were detected as cytoskeleton proteins in the NS-SV-AC spheroids (Supplementary Fig. 3 d & e, respectively).

In AGHA-3D culture spheroids, the protein location/profile expression (intensity detection) changed in most of the targeted proteins. AQP5 was translocated to the apical membrane in the majority of the cells within the spheroids, showing the characteristic profile for this protein (Fig. 5 i). The location was further confirmed by the z-stack 3D rendered image, where cells in the core of the spheroid also displayed APQ5 as transmembrane protein (Supplementary Fig. 4 a, Supplementary Movie 1). This protein was also detected in the cytoplasm but at a lower intensity. ZO-1 exhibited the characteristic distribution for tight junction proteins, where the connection between cells is point specific (Fig. 5 j); the 3D perspective showed an “*x,y,z*” cell-cell connection, confirming the junction in the three axes of the spheroid (Supplementary Fig. 4 b, Supplementary Movie 2).

I immunostaining indicated that the membrane co-transporter protein NKCC-1 was present at the edges of the spheroids, suggesting a basolateral distribution. However, low amounts of cytoplasmic NKCC-1 expression were also seen in the spheroids (Fig. 5 k, Supplementary Fig. 4 c, Supplementary Movie 3). α-amylase was detected mainly in the cytoplasm as small granules formed near the edges of the spheroids (Fig. 5 l, Supplementary Fig. 4 d, Supplementary Movie 4); the expression of this protein increased in 3D when compared with 2D culturing. The high expressions and locations of AQP5, ZO-1, and NKCC-1, as well as the self-assembled acini-like structures, confirmed the acinar identity of the 3D spheroids.

Similar to 2D, PanCK was detected in the NS-SV-AC spheroids (Supplementary Fig. 3 f). The expression of Col-IV and Pan-Laminin, the two most important proteins making up the basement membrane^2^, was detected only in a few spheroids among the population (data not shown). The expression of these proteins in the spheroids increased when compared with 2D culture, exhibiting the characteristic distribution at the periphery of the organized spheroids (Supplementary Fig. 3 g and h, respectively).

Based on these observations, the 2D culturing of NS-SV-AC leads to a fibroblast-like morphology after a few passages (∼>8) and/or at high cell confluency (> 95%), making it challenging to maintain an SG differentiated acinar state. There was a significant diminution in the expression of Vimentin (fibroblast marker) in NS-SV-AC cells when cultures in 3D compared to 2D (Supplementary Fig. 3 i *versus* Supplementary Fig. 3 d, respectively), suggesting that AGHA hydrogels offered better retention of SG epithelial phenotype than conventional 2D cultures. F-actin distribution was also distinct in 3D, showing a robust cell peripheral expression rather than cytoskeleton distribution, which was observed to be interconnecting with other cells from adjacent spheroids (Supplementary Fig. 3 j).

### NS-SV-AC form functional spheroids and respond to agonist-induced stimuli

The secretory function of NS-SV-AC cells cultured in 2D and 3D was evaluated by their sympathetic stimulation with 50 μM isoprenaline for 2 h. α-amylase activity was not found in the culture medium (supernatant) (data not shown) when tested using a commercial amylase assay. Further measurement of the intracellular α-amylase activity showed no activity in NS-SV-AC cells cultured as 2D monolayers (Fig. 6 a, circles), whereas NS-SV-AC spheroids in 3D culture showed amylase activity (Fig. 6 a, squares). While there were no significant differences (p>0.05) between stimulated and unstimulated samples, there was a significant difference observed in the amylase activity between 2D *versus* 3D platforms (Fig. 6 a).

**Fig. 6.**
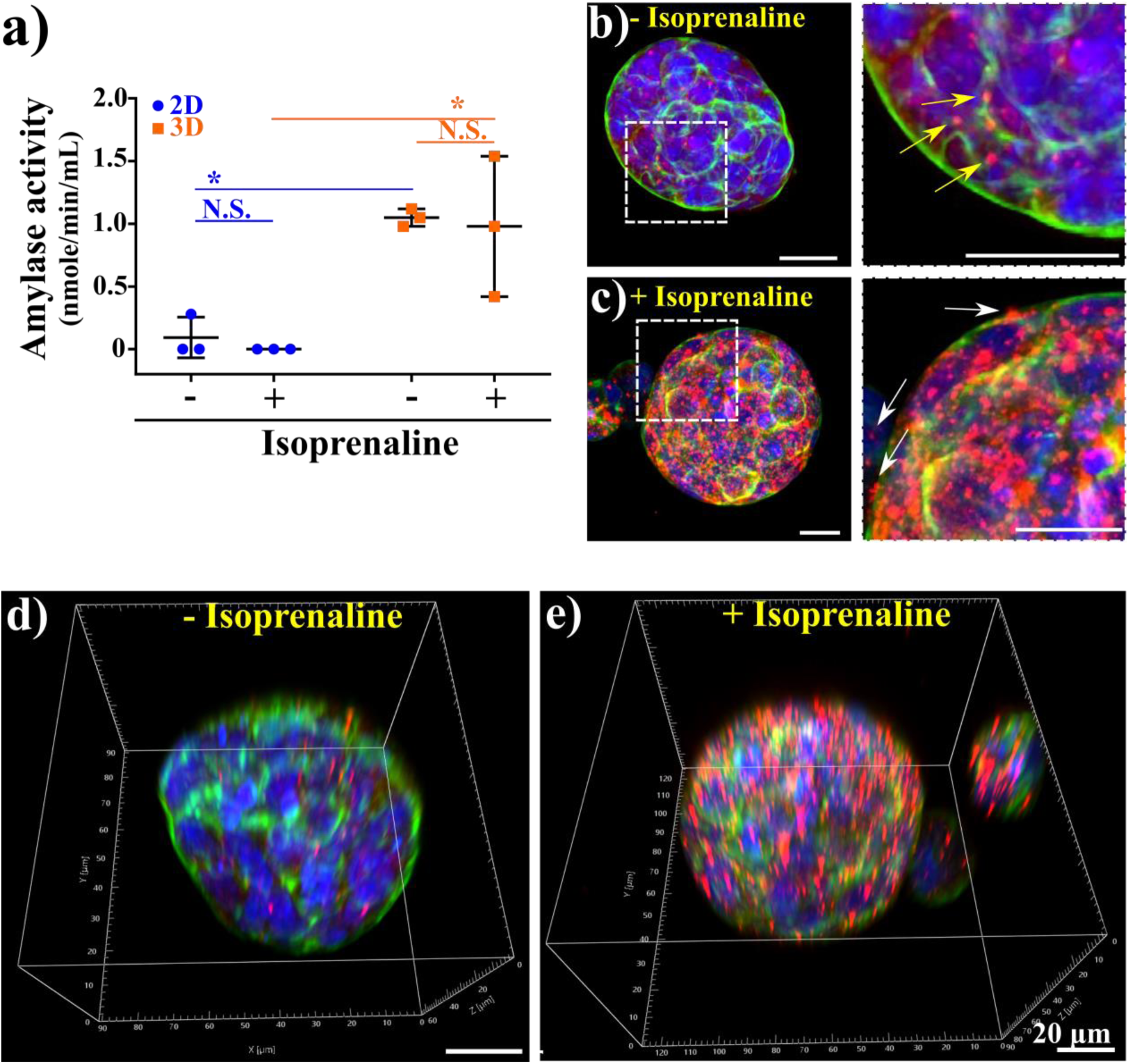
Functional analysis of NS-SV-AC spheroids cultured for 14 days in AGHA hydrogels. Cells and spheroids were stimulated with isoprenaline, and the amylase activity was measured by colorimetric assay. The amylase activity was reported as nmole/min/mL (milliunits), where one unit is the amount of amylase that cleaves ethylidene-pNP-G7 to generate 1.0 μmole of p-nitrophenol per minute at 25 °C. Data presented as mean ± SD, N.S. not significant, n≥3, *p<0.05. 2D maximum projection (b, c) and 3D reconstruction (d, e) of immunodetected α-amylase in non-stimulated (b, d) and stimulated (c, e). The panel at the very right is a close up of the selected area (white dotted box), and arrows indicate the α-amylase granules: red, α-amylase; green, F-actin; blue, nuclear staining. Magnification x20, scale bar 20 µm.

After treatment, 3D models were fixed and stained, then analyzed by confocal microscopy to determine the granule formation and α-amylase expression. Unstimulated spheroids produced a small number of α-amylase granules near the spheroid peripheries (Fig. 6 b, yellow arrows). Most of the immunodetected protein was distributed through the cytoplasm (Fig. 6 b), which can be considered as the basal levels of secretory granules and α-amylase staining. Post incubation with isoprenaline, there was an increase in intracellular secretory granule formation and a robust α-amylase staining were detected in all spheroids, with peripheral distribution and a few granules budding out from the cell membrane (Fig. 6 c, white arrows).

3D image reconstruction confirmed the spatial dispersion of α-amylase granules in unstimulated and stimulated samples, with granules close to cellular edges in the NS-SV-AC spheroids (delimited by F-actin staining) (Fig. 6 d, e, Supplementary Movie 5 and 6).

### AGHA hydrogels promote self-organization and maintenance of functional markers in 3D cultured human primary salivary cells

Considering the limited ability of cell lines to fully recapitulate native tissue characteristics, and as a proof of concept, we next sought to evaluate the performance of the AGHA hydrogels in culturing and expanding human-derived primary SG cells. Based on the previous studies, adult primary SG cells struggle to entirely reorganize into sizeable spheroids/organoids when cultured in dense matrices.^28^ For this, we used our previously well-characterized salivary functional units (SFU) model^4^ to evaluate the self-reorganization potential of SG cell clusters in 3D (Fig. 7 a).

**Fig. 7.**
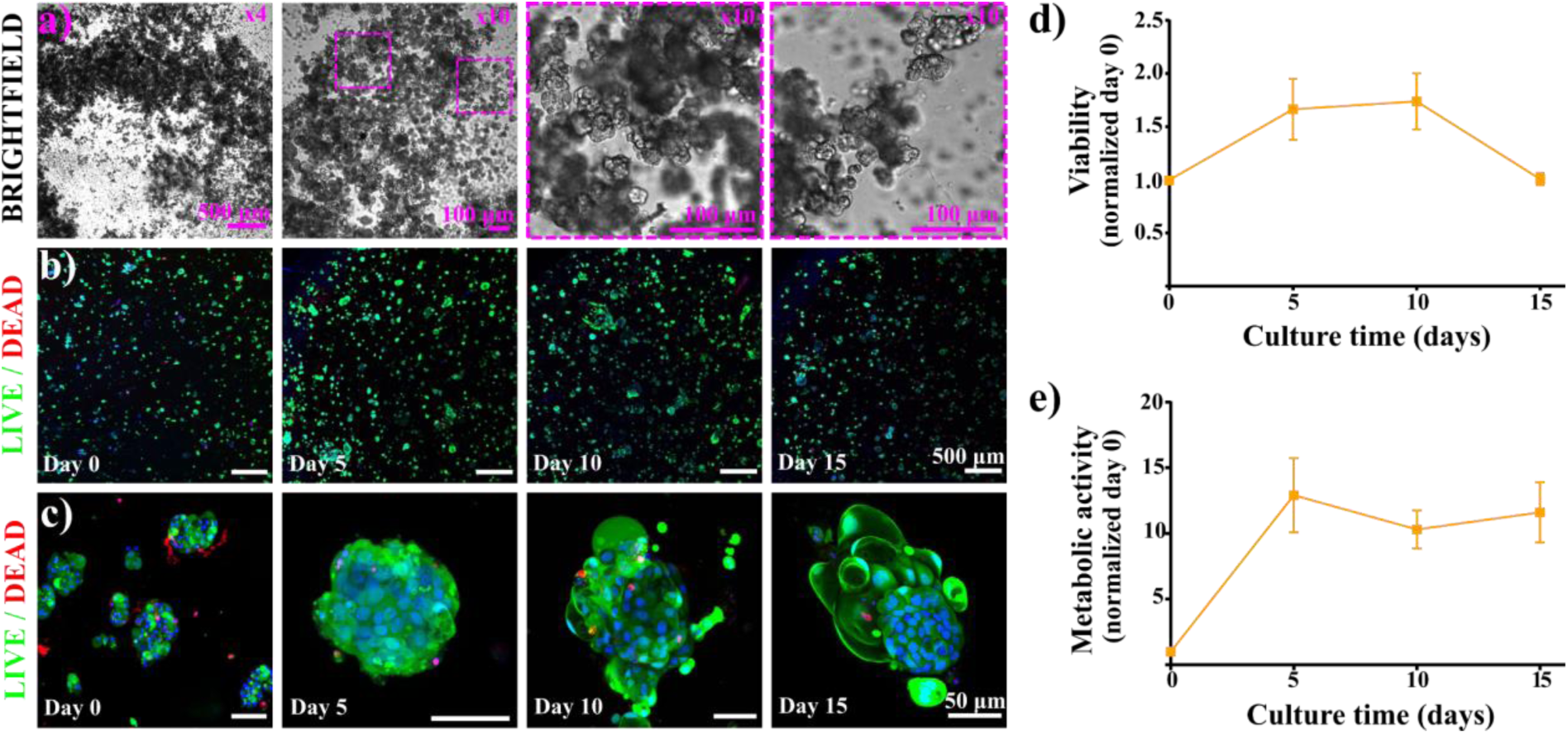
Isolation and 3D culture of human primary salivary cells (salivary functional units, SFU) in AGHA hydrogel system in culture for 15 days. a) Brightfield images of SFU isolated from fresh human salivary glands (day 0) at magnification of x4 and x10 (scale bar 500 μm and 100 μm, respectively). Confocal gallery of live/dead assay of SFUs culture in AGHA at a wide-field (b, x4/500 μm) and zoom-in (c, x20/50 μm) magnifications. SFU viability (d) and metabolic activity (e) were measured by live/dead and WST-1 assays, respectively. Green, live cells; red, dead cells; blue, nuclear staining. Data presented as mean ± SD, n≥3.

Live/dead assay revealed compact, self-organized SFUs to form sizeable SG microtissues while maintaining high levels of viability over 10 days, which then, decreased at day 15 (Fig. 7 b-d). Metabolic activity of 3D cultured SFUs showed an exponential increase in metabolic activity within the first 5 days of culture, followed by a plateau phase sustained up to day 15 (Fig. 7 e). The metabolic spurt in the SG microtissues during the first five days was concurrent with our previous findings when SFUs were grown in suspension cultures.^4^ Using our biomimetic hyaluronic acid-based 3D matrix (AGHA gel), we report an improved 3D culture platform for the long-term sustenance of actively aggregating, viable SFUs towards organoid formation.

Human SFUs showed the high expression and proper location of acinar markers such as AQP5, ZO-1, NKCC1, and α-amylase (Fig. 8 a-d, Supplementary Fig. 5 a-d, Supplementary Movies 7-10). Human SFUs exhibited the expression of Epithelial Cadherin (ECad was used here as a marker for adherens junctions; Fig. 8 a, Supplementary Fig. 5 a, Supplementary Movie 7). Vimentin, Col IV, and Pan-Laminin were detected only in a few cells contained within the SFUs (Fig. 8 e-g). F-Actin staining revealed that the well-compact cell organization of SFUs was maintained (Fig. 8 h). Given together, these results suggest that AGHA gels promoted and maintained the differentiated acinar state of the human primary SG cells for at least 15 days.

**Fig. 8.**
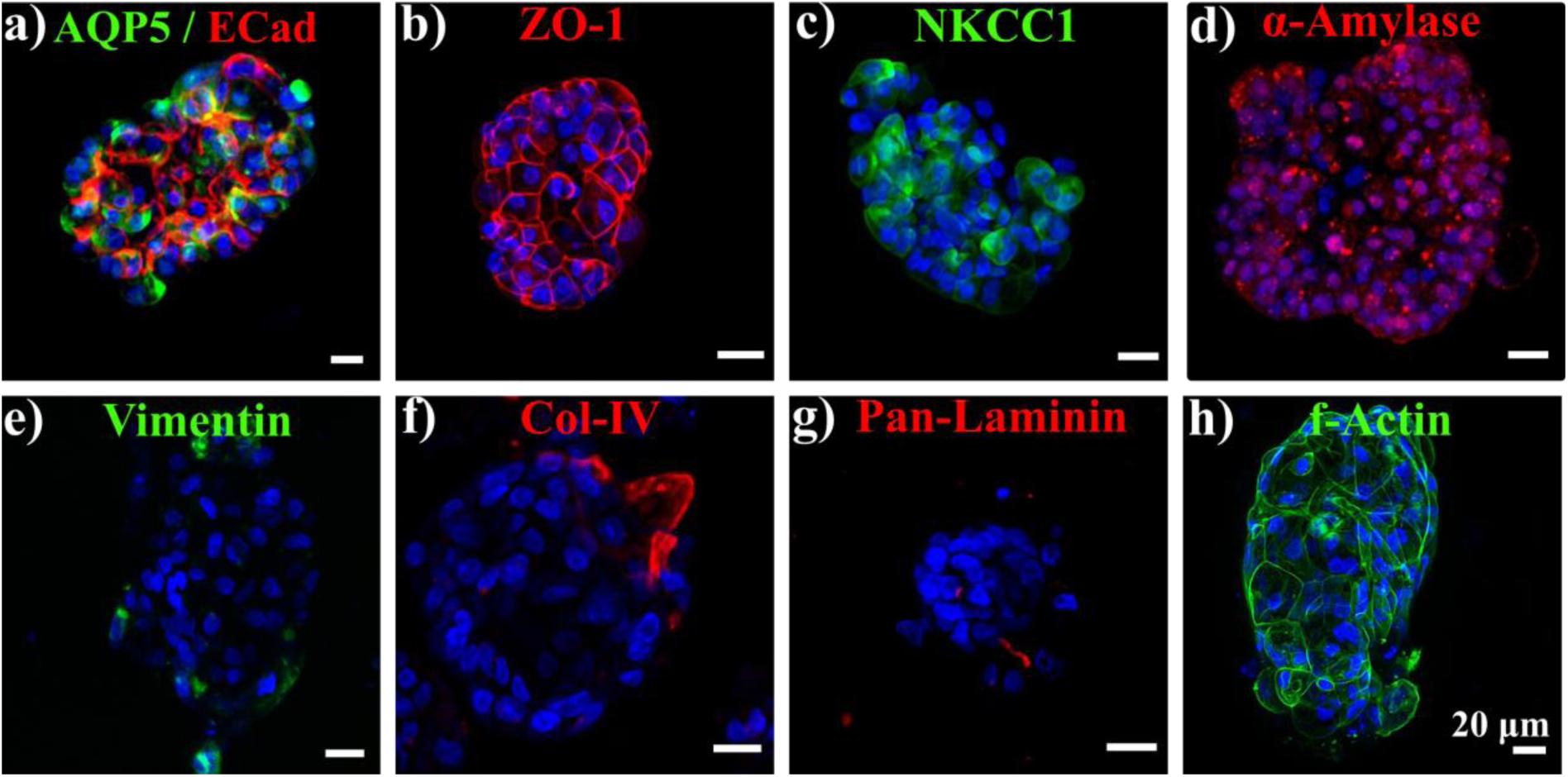
Immunodetection of SG markers in human SFU expanded in 3D AGHA hydrogels cultured for 15 days. 2D maximum projection confocal images of SG markers expressed by SFU (a-d). Green, AQP5, NKCC1, Vimentin, or f-actin; red, Ecad, ZO-1, α-Amylase, Col-IV, or Pan-Laminin; blue, nuclear staining. Magnification x20, scale bar 20 µm.

## Discussion

Although different platforms were developed in the field of SG bioengineering to recapitulate the 3D environment of salivary cells *in vitro*, these 3D models and culture conditions are still being optimized. It remains challenging to maintain a differentiated acinar state and cellular heterogeneity^29^, especially using primary cultures. Due to the shortage of SG patient biopsies, different approaches have been followed to ensure enough cell concentration to study SG biology *in vitro*, such as tissue and slice culture, cell isolation and expansion in 2D, and recently, the use of 3D culture models that recapitulate the native environment. Immortalized cell lines are helpful in standardizing and optimizing new *in vitro* and *ex vivo* 3D biomaterial-based disease models. We used the human salivary cell line (NS-SV-AC) and human primary salivary cells (SFU) as biological models to test our SG 3D systems.

Alginate-based hydrogels have been extensively used as 3D scaffolds to grow cancer^8–10,13,30^ and non-cancer cells,^31,32^ including SG cells.^33–35^ Recently, HA hydrogels have been proposed as a great niche for SG spheroids.^23^ Nonetheless, the addition of collagen or HA into AG hydrogels as freely suspended moieties to create stable, reversible 3D matrices has been poorly explored. We developed and tested three different hydrogel blend compositions to expand and promote the formation of human SG acinar spheroids in 3D while maintaining their acinar-differentiated state. It has been reported that incorporating other biopolymers to AG gels, such as collagen or HA, besides providing biological cues to the matrix, can either increase or reduce its stiffness, which can also modulate the behavior of seeded cells and contribute to the formation of spheroids.^10,13^ The mechanical characterization of our thermal-ionic crosslinked gels did not show significant differences, and most importantly, the values obtained from ElastoSens Bio2 (viscoelastic properties, Fig. 2) and micro-compression (Fig. 3) tests revealed comparable mechanical properties to those reported for human SG tissues (9-33 kPa).^36^ We hypothesize that thermal-ionic crosslinked methods to form these hydrogels could generate a two-phase 3D model, comprised of a partially-crosslinked softcore surrounded by a stiffer shell. The addition of collagen or HA into AG gels might be interacting with alginate chains altering the “egg-box” formation, ergo, modifying the crosslinking speed (outside and inside of the alginate), network, and final stiffness.^37,38^

Although all tested hydrogels enable the self-assembly of NS-SV-AC in well-organized, compact 3D structures, resembling the basic acini units in SG structures,^2,39^ the matrix composition highly influenced its viability and metabolic activity. HA-based hydrogels have been used previously as 3D platforms to expand and produce functional SG spheroids, in particular, using covalent cross-linking reactions to form stable hydrogels^2,18,19,40^ or by immobilizing the HA to a material surface,^23^ requiring a long enzymatic digestion process (∼4 h) for further spheroid isolation, which may affect the cell state of the retrieved cells due to the extended exposure to dissociative enzymes.^41^ In our 3D system, we used temperature (25 °C) and calcium ions as a crosslinker to form the 3D matrix, representing an advantage in cell/spheroid recovery by a simple ion chelation process (<5 min) at 37°C for downstream applications, avoiding the aggressive enzymatic treatment while keeping spheroid integrity.

The presence of HA in our 3D AGHA hydrogel could be one of the main reasons for the matrix’s preference for human SG acinar cells since this cell type (acinar cells e.g., NS-SV-AC) contain CD44 receptors,^20,42^ which are the HA binding-receptors^21,22^ interacting and activating essential pathways to successfully promote the formation of acini structure in *in vitro* systems. AQP5 is a water channel protein and is one of the most important proteins involved in saliva production by SG acinar cells. This protein is localized in cytoplasmic vesicles that can be translocated to the cell apical plasma membrane in response to stimuli for saliva secretion. The proper expression and translocation of AQP5 is essential for saliva production.^43^ NS-SV-AC spheroids produced in our AGHA system expressed AQP5 as a transmembrane protein in the apical side of the cell membrane, which suggests a proper protein translocation to ensure its correct functionality. The tight junction protein ZO-1 showed its characteristic distribution as a point-specific interconnecting protein between cells. In epithelial tissues, the establishment of cell polarity is closely linked to the establishment of the apical junction complex, which includes tight and adherent junction proteins, creating a barrier preventing lateral movement of proteins between apical and basolateral membranes^44,45^. The presence of ZO-1 suggests NS-SV-AC cell polarity in the spheroids.^46,47^ Furthermore, the membrane co-transporter protein NKCC-1 exhibited a basolateral distribution in the cells, with similar patterns to the one reported by other authors.^48–50^ These results demonstrated that salivary spheroids cultured in our hydrogel maintained robust expression of key proteins that remained correctly localized on cell membranes.

Salivary α-amylase, commonly used as a marker of secretory function in the SG,^48^ is one of the most abundant components in saliva locally produced by highly differentiated acinar cells, mainly from the parotid glands^51^ and secreted in response to stimulation of adrenergic receptors in these cells.^52^ Our 3D acinar spheroids showed higher basal cytoplasmic expression of α-amylase compared with 2D culture under non-stimulated conditions. These results confirmed the acinar identity of the 3D spheroids. The fact that 2D cultured acinar cells presented a lower expression and altered location suggests a loss of the differentiated acinar-like state in these cells, which was somehow restored in our 3D culture system.

When the secretory function of NS-SV-AC after sympathetic stimuli was evaluated, we did not find α-amylase activity in the supernatant but expressed intracellularly, showing higher activity in 3D cultures compared to 2D systems. The secretory process in exocrine glands (such as SG) involves sequential steps such as secretory vesicle trafficking, tethering, docking, priming, and membrane fusion; interference in any of these steps can lead to altered secretion of salivary proteins^53^. We hypothesize that 3D acinar spheroids are needed from the participation of additional cell types enabling the full secretion of α-amylase to the medium. These co-culture assays must be performed to confirm our hypothesis.

In the AGHA-3D spheroids, we confirmed the robust α-amylase expression with an increment in the intracellular secretory granules post-stimulation with isoprenaline, where the α-amylase exhibited peripheral distribution and granules budding out from the cell membrane, indicative of exocytosing vesicles.^19^ While it has been shown that basal amylase activity on primary SG stem cells seeded in decellularized ECM from the submandibular gland showed 14 times higher activity than 2D culture platforms after five days of culture, the study did not extend beyond this time point.^54^ It has also been reported that α-amylase activity decreased over time in SG tissue (slice culture) and primary SG cell culture, losing up to 90% of its activity after 14 days of culture^3^. However, several studies using isoprenaline to stimulate the SG (PG, SMG) mainly report the immunodetection (Western blot, immunofluorescence)^48,55^ or enzymatic activity^52,55^ of amylase. Just a few ones have shown both assays, but in 2D system,^52,56^ the formations of amylase secretory granules are not clearly defined.^57^ Our stimulated acini spheroids showed functionality and high expression of α-amylase, with a peripheral distribution and granules budding out from the 3D acini structure.

## Conclusions

Overall, the three hydrogels tested show mechanical properties within the range of human SG. Although NS-SV-AC cells can self-assemble into spheroids in all three hydrogels, the AGHA hydrogel provides the best mechanical and biochemical properties for SG cell expansion and spheroid formation, promoting the production of larger, viable, and well-organized 3D structures. This 3D *in vitro* models benefits the proper location and high expression of key acinar cell markers, producing functional human SG acinar spheroids. Our HA-based platform also represents an advantage as a cell expansion system due to the reversibility of the 3D matrix formed, allowing the easy recovery of the produced spheroids by a simple ion chelation process at 37°C without compromising their integrity, viability, and functionality for downstream applications such as single-cell isolation and scale up spheroid production. For this work, we primarily optimized the more sensitive and difficult-to-culture acinar cells which are the main secretory cells in SGs. Most importantly, our 3D-AGHA models support the expansion of human SFU (primary salivary cells) while maintaining high viability and expression of key SG markers. This HA-based platform has the potential to be used as *in vitro* and *ex vivo* 3D disease modeling platform (for example, radiation response model) to expand diverse SGs cellular populations such as the ductal, myoepithelial, and mesenchymal cells. These heterogenous spheroid populations can be used in long-term co-culture systems to integrate a complex *ex vivo* salivary gland architecture.

## Materials and Methods

### Preparation of hydrogels

For alginate/gelatin (AG) hydrogels, 1% (w/v) sodium alginate (Protanal LF 10/60 FT, FMC BioPolymer) and 7% (w/v) gelatin from bovine skin Type B (Sigma) sterile powder were dispersed into Dulbecco’s phosphate buffer saline (DPBS, without calcium and magnesium, Wisent BioProducts) by mixing for 3 h at 50 °C/500 rpm with a magnetic stirrer. Hydrogel solids were dissolved (sol-liquid transition phase) at 37 °C for 30 min before being transferred into clean, sterile 50 mL conical tubes (Corning) inside of a biological safety cabinet (BSC). For alginate/gelatin/collagen (AGC) hydrogels, bovine collagen type I (5 mg/mL, Gibco) was added to 30 °C pre-warmed A1G7 solutions at a final concentration of 5% (v/v) and mixed for 30 min with a magnetic stirrer at 30 °C. All steps were performed under sterile conditions.

For alginate/gelatin/hyaluronic acid (AGHA) hydrogels, 10 mg of HA vial (Glycosil thiol-modified hyaluronan, Advanced Biomatrix) was dissolved into 1 mL of degassed, deionized, sterile water. Then, 7.5% (v/v) HA was added into a 30 °C pre-warmed AG solution and mixed for 30 min at 30 °C. Alginate and gelatin powders were sterilized under UV light for 12 h and weighed inside a BSC. All hydrogels were centrifugated at 2,000 rpm for 10 min to eliminate gas bubbles and transferred into sterile 10 mL syringes. Hydrogel preparation was carried out under sterile conditions within a BSC and stored at 4 °C until use.

### Hydrogel mechanical testing

Viscoelastic properties of hydrogels were analyzed using a non-invasive and contact-free mechanical tester ElastoSens^TM^ Bio2 (Rheolution Inc, Canada) to measure the storage modulus (G’) and micro-compression tester (Microsquisher, CellScale Biomaterials, Waterloo, ON, Canada) for compression modulus (see Supplementary Information for details).

### Cell culture

For optimization of the 3D models, we used the human cell line Normal Salivary Simian Virus 40-immortalized Acinar Cells (NS-SV-AC) for the biological validation of our gels and culture conditions. The cells were negative to mycoplasma. The NS-SV-AC were cultured in complete Mammary Epithelia Growth media (Epi Max I, Wisent BioProducts), plus 100 U/mL penicillin, 100 μg/mL streptomycin, 0.25 μg/mL amphotericin B (Gibco) and 10% fetal bovine serum (FBS) at 37 °C and 5% of CO_2_ until 80% confluence. Cells were harvested by incubation with a trypsin-EDTA (Gibco) solution and transferred into a new, sterile culture dish (Corning). All cells were used from passages 3 to 6 after thawing from liquid nitrogen storage. The medium was changed every two days as follows: all culture medium was recovered and filtered through sterile filters of 0.22 µm pore size(conditioned medium); then, cells were washed with DPBS, and culture medium was added at a 2:1 ratio (2 parts of fresh complete Epi Max I medium plus 1 part of conditioned medium). Culture dishes were incubated using the above conditions.

As a proof of concept and to validate our best 3D candidate, we isolated and cultured human primary salivary cells (that we named SFU) and seeded within our AGHA gel. Human salivary glands were obtained from consenting patients undergoing surgery for various diseases and obtained according to the McGill Institutional Review Board guidelines (IRB no: A05-M62-05B). Freshly isolated salivary glands (∼3-5 g) were processed as per our previously established guidelines (see Supplementary Information for details) ^4,58^.

### Viability, size of spheroids, and metabolic activity

Hydrogels were incubated for 30 min at 37 °C in a water bath to allow the sol-gel transition. In parallel, NS-SV-AC cells were harvested, and 2x10^6^ cells/mL were mixed into 1 mL of hydrogels. Thirty-microliter hydrogel drops (3D models) were created by the hanging drop method on the surface of a 100 mm sterile culture dish, and the 3D models were left to settle for 5 min to thermally cross-link the gelatin. Then, sterile 100 mM CaCl_2_ solution was added to the top of the 3D models for ionic cross-linking and incubated for 2 min at RT. 3D models were carefully liftoff from the culture dish surface with a spatula to allow the complete cross-linking of the 3D models. The 3D models were rinsed with DPBS and transferred into an agarose-coated 6-well plate. Complete Epi Max I culture medium was added to each well and cultured at 37 °C for 14 days, depending on the experiment. The medium was changed every three days following the methodology above.

Cell viability was determined using Calcein-AM (AAT Bioquest) and Ethidium homodimer I (Biotium) (Live/Dead assay); the working solution was prepared by diluting 1 µl of Calcein-AM (4 mM) and 2 µl of Ethidium homodimer I (2 mM) into 1 mL of DPBS. As a nuclear stain, two µL of Hoechst 33342 (18 mM, TOCRIS Bioscience) was added to the working solution. Samples were incubated at 37 °C for 45 min, and the 3D models were rinsed twice with DPBS at RT. Confocal images were acquired using a Nikon A1 laser confocal with a Z-stack scan of 100-200 µm thickness and 1-1.5 µm steps at magnifications of ×10 and ×20 and viability and size of spheroids were calculated on processed images (see Supplementary Information for details).

The metabolic activity (proliferation) of spheroids was measured using Cell Proliferation Reagent WST-1 (Roche®), following the manufacturer’s instructions. Briefly, 3D models were rinsed with DPBS, and 100 µl of fresh culture medium plus 10 µl of WST-1 Reagent were added to each sample following an incubation at 37 °C for 2 h. Then, the volume was transferred into a new, clean 96-well plate, and absorbances were measured at 440 nm in a nanodrop spectrophotometer (NanoDrop 2000, Thermo Scientific).

### Quantitative real-time polymerase chain reaction

The levels of transcripts of SG markers of NS-SV-AC cultured in 2D (4 days in culture) and 3D AGHA (14 days in culture) were determined by real-time polymerase chain reaction (qPCR).

The detailed process and primer sequence (Supplementary Table 1) can be consulted in the Supplementary Information.

### Whole-mount staining for SG markers expressed by 3D NS-SV-AC spheroids

The 3D models were collected from the culture and rinsed twice with NaCl/HEPES buffer (135 mM NaCl, 20 mM HEPES, pH 7.4).^59^ Immunostaining and confocal imaging were performed to detect the SG marker expressed by the 3D NS-SV-AC spheroids. The antibodies (Supplementary Table 2) and the detailed protocol can be consulted in Supplementary Information. NS-SV-AC spheroids were released from the gel for better confocal image acquisition. Imaris Viewer 9.9.0 software was used to render 3D perspective images on processed confocal images.

### Functional analysis of 3D NS-SV-AC spheroids by amylase activity

The secretory function of NS-SV-AC culture in 2D and 3D platforms were evaluated by stimulating cells with sympathetic stimuli. Cells were cultured in 2D platforms at 90-95% confluency (4 days in culture); the cells were stimulated with 50 µM isoprenaline (Sigma Aldrich) for 2 hours at 37 °C/5% CO_2_ and harvested. Four million cells were mixed with amylase lysis buffer and homogenized. The amylase activity was determined using a colorimetric assay kit (MAK009, Sigma-Aldrich) and ethylidene-pNP-G7 as substrate following the manufacturer’s instructions with a modified protocol by employing the whole cell extract rather than the supernatant. Samples were measured at 405 nm using a microplate reader (EL800, Bio-Tek Instruments). Culture without isoprenaline stimulation was used as a control. For the 3D platforms, first, the number of cells per 3D model was estimated. The spheroids and cells from five 3D AGHA models (14 days in culture) were released from the gel and incubated with trypsin for 2 min at 37 °C. Then, samples were centrifuged at 1,000 rpm, and the pellet was resuspended in culture medium. The cell concentration was determined by trypan blue assay, and the number of cells per 3D model was calculated by dividing the total cell amount by the number of dissociated gels.

Thirty 3D models (∼4x10^6^ viable cells, 14 days of culture) were incubated with 50 mM isoprenaline for 2 hours at 37 °C/5% CO_2_. After stimulation, the 3D models were collected, and the spheroids were released from the gel, as explained earlier. The amylase activity was determined following the same procedure as for 2D culture. Amylase activity was reported as nmole/min/mL (milliunits), where one unit is the amount of amylase that cleaves ethylidene-pNP-G7 to generate 1.0 μmole of *p*-nitrophenol per minute at 25 °C. Thirty hydrogels were placed into plain medium as control. The experiments were performed in triplicates.

### Statistical Analysis

All experiments were performed as quadruplicated unless stated otherwise. For viability and metabolic activity tests, data were normalized to day 0 values; for qPCR, data were normalized to GAPDH values. Data are presented as mean ± SD. *t*-test and One- or Two-way ANOVA was performed with Tukey’s, Dunnett’s, or Bonferroni’s post hoc test with a p-value <0.05. Data were plotted using GraphPad Prism 9.0. Surface area values were plotted as violin plots using web-based tools PlotsOfData and PlotsOfDifferences.^60^

## Acknowledgements

We thank the financial support from Fonds de Recherche du Québec Santé (FRQS, grant no. 281271). We would like to thank Dr. Masayuki Azuma for providing the NS-SV-AC cell line, Dr. Younan Liu for his technical assistance, and Dr. Dieter Reinhardt and Dr. Christine Delporte for their comments on some experiments. SP acknowledges support from FRQS doctoral award #304367. SNN acknowledges funding from CFI, Rheolution Inc., and Investissement Québec.

## Contributions

**JGML:** Conceptualization, methodology, validation, formal analysis, investigation, writing-original draft, visualization, project administration, writing-review & editing. **SP:** Conceptualization, methodology, validation, formal analysis, investigation, writing-original draft, visualization, writing-review & editing. **YZ:** Methodology (metabolic assay), validation (metabolic assay) writing-review & editing. **AG:** Methodology (metabolic assay), validation (metabolic assay), writing-review & editing. **DC:** Methodology (mechanical testing), writing-review & editing. **SNN:** Resources (mechanical testing), writing-review & editing. **JMK:** Resources, writing-review & editing, supervision, funding acquisition. **SDT:** Resources, writing-review & editing, supervision, project administration, funding acquisition.

## Supplementary data

Supplementary data to this article can be found online at.

