## Supplementary Information for "Expansion of functional human salivary acinar cell spheroids with reversible thermo-ionically crosslinked 3D hydrogels"

### Materials and Methods

#### *Hydrogel mechanical testing*

To measure the storage modulus (G') values achieved by gelatin gelation (themogelation), hydrogels were pre-warmed at 37 °C, and seven mL per sample were loaded into a holder located inside of the ElastoSens$^{TM}$ Bio2 and the data acquisition started immediately. The principle of the instrument is to apply a low frequency-amplitude (gentle mechanical) vibration to the rigid holder, which has a flexible, bottom silicone rubber membrane; then, the dynamic response of the sample is measured using a contactless laser probe and processed utilizing specialized models to obtain the shear storage modulus (G') of the sample.[1] The tests were performed at 25 °C for 180 min, with a temporal step of 30 s using the "soft mode" parameters. Data were plotted as a function of time, and the values from the last 5 min were used to determine differences in the maximum average G' of hydrogels. To measure the G' values obtained by ionically crosslinked gels, three mL of gel were loaded into a holder and incubated for 10 min at 4 °C to speed up the gelation of the gelatin; then, three mL of 100 mM $CaCl_2$ solution were added to the top of the gels and incubated at 25 °C for 30 min. The tests were performed at 25 °C for 4 min, with a temporal step of 5 s using the "stiff mode" parameters. Data were plotted as a function of time, and the values from the last minute were used to determine differences in the maximum average G' of hydrogels. For compression testing, 30 µL hydrogel drops (3D models) were created by the hanging drop method on the surface of a 100 mm sterile culture dish, and the 3D models were left to settle for 4 min at RT followed by 1 min at 4 °C to thermogelation of the gelatin. For thermos-ionically crosslinked gels, sterile 100 mM $CaCl_2$ solution was added to the top of the 3D models and incubated for 5 min at RT. Then, 3D models were lifted off from the culture dish surface with a spatula to allow the complete cross-linking of the 3D models. Samples were rinsed with DPBS. Both batches, thermo and thermo-ionically crosslinked, were gently cut using a ø3 mm biopsy punch to create cylindrical specimens. The parallel plate compression was performed using a micro compression tester (Microsquisher, CellScale Biomaterials, Waterloo, ON, Canada) coupled to a ø 0.5588 microbeam with 6x6 mm platen and with a displacement rate of 30 µm/s. Stress-strain curves were generated using the "force *versus* displacement" data given by the instrument and the heights and initial cross-sectional areas of the hydrogels. The stress-strain curves were plotted from 0 to 0.35. Slopes of the linear portion of these stress-strain curves (0.05 to 0.10) were used to calculate the compressive modulus.

*Cell culture: isolation of salivary functional units (SFU)*

Freshly isolated human salivary glands (~3-5 g) glands were washed thrice using phosphate buffer saline (PBS) containing 100 U/mL penicillin, 100 µg/mL streptomycin, and 0.25 µg/mL amphotericin B. Then, the connective tissues and surrounding non-glandular structures were manually removed. The gland tissue was sectioned into small pieces using a scalpel, followed by scissors cutting to homogenize the tissue (slurry). The homogenized tissue was transferred into a 50 mL clean, sterile conical tube containing 35 µg/mL Liberase enzyme (Roche®) in MEM (supplemented with 100 U/mL penicillin, 100 µg/mL streptomycin, and 0.25 µg/mL amphotericin B). Then, the sample was briefly dissociated using a GentleMACS (Miltenyi biotech) dissociator followed by incubation at 37°C and 5% $CO_2$ with manual shaking every 30 mins until all tissues are visibly digested. Digested glands were centrifuged at 1,500 rpm for 5 mins. The pellet was carefully resuspended in PBS and sequentially passed through a 70 µm and a 20 µm cell strainer. SFU (ø70 µm-20 µm) were collected and washed three times with 0.9% normal saline to remove debris and SFU were cultured in AGHA hydrogels as discussed above using complete Epimax supplemented with 10% FBS.

*Viability, size of spheroids, and metabolic activity*

Cell viability was determined using Live/Dead assay and confocal images were acquired using a Nikon A1 laser confocal with a Z-stack scan of 100-200 µm thickness and 1-1.5 µm steps at magnifications of ×10 and ×20. The noise reduction and processing of the acquired images were performed with Fiji software.[2-4] "Median 3D", "unsharp mask," and "remove outliers" filters were used as the first cleaning step. Using a maximum stack arithmetic tool ("Z projection" plug-in), the 2D image projections were created, the background removed based on the "rolling ball" algorithm (50-500 pixels depending on the spheroid size) and adjusting brightness and contrasts. The fluorescence intensity obtained from processed Z-stack confocal imaging was used as a parameter to determine the viability of spheroids/cells in culture. On the other hand, the spheroid surface area was used as a parameter to determine the quantity and sizes of the spheroids produced into hydrogels; sizes larger than 400 µm$^2$ were considered as spheroids.[5-7] Manual segmentation and selection of spheroids were used to delimit the surface area of each spheroid.

The metabolic activity (proliferation) of spheroids was measured using Cell Proliferation Reagent WST-1 (Roche®), following the manufacturer's instructions. 3D models were rinsed three times with DPBS, and 100 µl of fresh culture medium plus 10 µl of WST-1 Reagent were added to each well. Samples were incubated at 37 °C for 2 h. Then, the volume was carefully transferred into a new, clean 96-well plate, and absorbances were measured at 440 nm in a nanodrop spectrophotometer (NanoDrop 2000, Thermo Scientific).

*Quantitative real-time polymerase chain reaction*

The levels of transcripts of SG markers of NS-SV-AC cultured in 2D (4 days in culture) and 3D AGHA (14 days in culture) were determined by real-time polymerase chain reaction (qPCR). For 3D samples, first, acinar spheroids were released from gels by ion chelation process.[8] The hydrogels were transferred into clean 1.5 mL Eppendorf tubes, and 100 µL (per 3D models) of pre-warmed (37°C) 55 mM trisodium citrate (Sigma) were added. Hydrogels were fully dissociated by gently pipetting for one minute, and samples were centrifuged at 1,000 rpm/ 5 min /RT. The supernatant was removed, and spheroids were rinsed with DPBS thrice, repeating the centrifugation step between washes. Total RNA was isolated using RNeasy Plus Mini Kit (Qiagen, Canada), and the concentration was measured at 260 nm absorbance using a spectrophotometer (SmartSpec 3000, Bio-Rad). The complementary DNA (cDNA) was synthesized using 1 µg of mRNA and a high-capacity cDNA reverse transcription kit (Applied Biosystems, Canada) employing a gradient 96 well DNA engine thermal cycler (Bio-Rad, Canada) according to the manufacturer's instruction. The qPCR reaction was set up using 10 µl 2X SYBR Green master mix, 2 µl of cDNA, 1 µl of each forward and reverse primers (100 nM), and 6 µl of nuclease-free water to a final reaction volume of 20 µl and the reaction was run using StepOnePlus real-time PCR (Applied Biosystems, Canada). The primer sequences used are listed in Supplementary Table 1. All qPCR reactions were performed as biological replicates in three independent experiments, and each gene was compared with the housekeeping gene GAPDH. The relative gene expression was expressed as fold change and calculated using the $2^{-\Delta\Delta Ct}$ method.[9]

*Whole-mount staining for SG markers expressed by 3D NS-SV-AC spheroids*

The 3D models were collected from the culture and rinsed twice with NaCl/HEPES buffer (135 mM NaCl, 20 mM HEPES, pH 7.4). First, gels were cross-linked again by incubating into 100 mM $CaCl_2$ solution for 1 min and rinsed once with NaCl/HEPES buffer. Then, the 3D models were fixed with 4% paraformaldehyde in NaCl/HEPES buffer for 1 h at RT. Samples were washed with NaCl/HEPES buffer thrice for 15 min each. Unspecific proteins were blocked with a blocking solution (5% BSA in NaCl/HEPES buffer) for 4 h at RT. After blocking, hydrogels were cross-linked (as described above) to ensure no gel dissociation during the subsequent steps. The antibodies used are listed in Supplementary Table 2: The primary antibodies were diluted in 3% BSA in NaCl/HEPES and incubated overnight at 4 °C. Samples were washed with NaCl/HEPES buffer thrice for 15 min each. A third cross-linking incubation is required before the final steps. Then, the secondary antibodies were diluted in 3% BSA in NaCl/HEPES and incubated overnight at 4 °C. The samples were washed with NaCl/HEPES buffer thrice for 15 min each and stained with Hoechst 33342 (18 mM) (3 µl per one mL of NaCl/HEPES solution) for 20 min at RT.

**Table 1**: Primers used for qPCR

| Target gene | | Primer sequences (5’-3’) |
|---|---|---|
| **AMY-1** | F | CCTTCTGGGATGCTAGGCTG |
| | R | ATCTTGGCCAACGGTAGCTT |
| **AQP-5** | F | GCTCACTGGGTTTTCTGGGTA |
| | R | CCTCGTCAGGCTCATACGTG |
| **GAPDH** | F | AGGGCTGCTTTTAACTCTGGT |
| | R | CCCCACTTGATTTTGGAGGGA |
| **NKCC-1** | F | CCTCTACACAAGCCCTGACTTAC |
| | R | CGTGAGTTTGGAGCACCTGTCA |
| **ZO-1** | F | CGGTCCTCTGAGCCTGTAAG |
| | R | GGATCTACATGCGACGACAA |

AMY-1: amylase; AQP-5: aquaporin-5; GAPDH: glyceraldehyde-3-phosphate dehydrogenase; NKCC-1: sodium-potassium-chloride channel; ZO-1: zona occludens-1. F: forward; R: reverse

**Table 2.** List of antibodies used for immunodetection.

| Antibody | Acronym | Working dilution | Brand | Catalog number |
|---|---|---|---|---|
| **Primary** | | | | |
| Rabbit polyclonal anti-aquaporin-5 | AQP5 | 1:125 | Thermofisher | PA5-99403 |
| Recombinant rabbit monoclonal anti-Zona Occludens-1, EPR19945-296 | ZO-1 | 1:100 | Abcam | ab221547 |
| Rabbit polyclonal wide spectrum anti-cytokeratin | PanCK | 1:125 | Abcam | ab9377 |
| Rabbit polyclonal anti-collagen IV | Col-IV | 1:200 | Abcam | ab6586 |
| Rabbit polyclonal anti-laminin | Pan-Laminin | 1:100 | Thermofisher | PA1-16730 |
| Recombinant Alexa Fluor 488 anti-vimentin, EPR3776 | Vimentin | 1:200 | Abcam | ab185030 |
| Recombinant anti-salivary alpha-amylase | α-Amylase | 1:125 | Abcam | ab201450 |
| Mouse monoclonal anti-E-cadherin | E-Cad | 1:125 | Abcam | ab1416 |
| Phalloidin-iFlour 488 conjugate | F-actin | 1:500 | AAT Bioquest | 23115 |
| **Secondary** | | | | |
| Goat anti-Rabbit IgG (H+L) Cross-Adsorbed Alexa Fluor 594 | N.A. | 1:200 | Thermofisher | A-11005 |
| Goat anti-Mouse IgG (H+L) Cross-Adsorbed Alexa Fluor 594 | N.A. | 1:200 | Thermofisher | A-11012 |

N.A.; Not applicable.

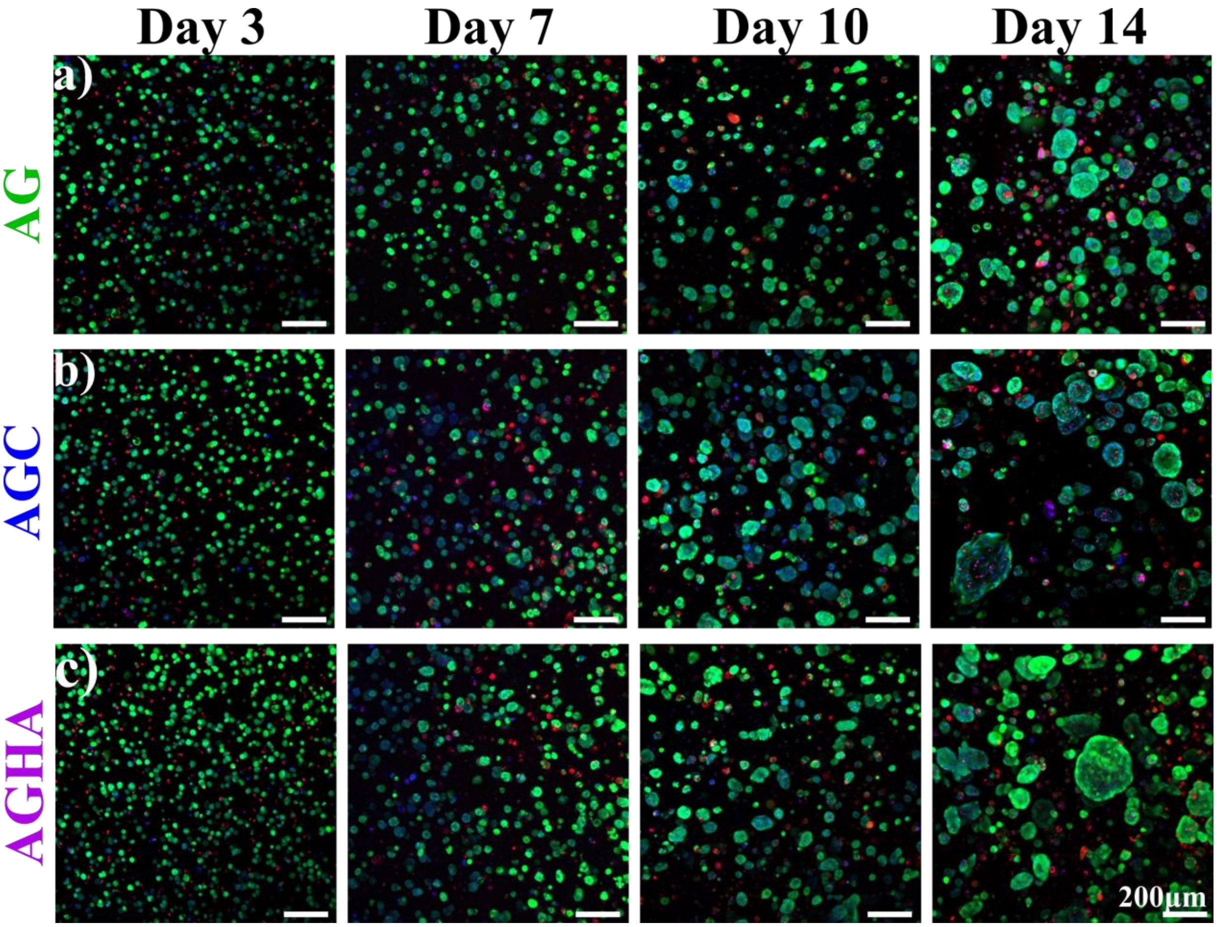


**Fig. 1**. NS-SV-AC spheroid formation in 3D hydrogels. Cells were seeded into a) AG, b) AGC, or c) AGHA and cultured for 14 days. Live/dead assay confocal imaging was performed to determine the spheroids' viability, morphology, and size. Green, live cells; red, dead cells; blue, nuclear staining. Magnification/scale bar: x10/200 μm.

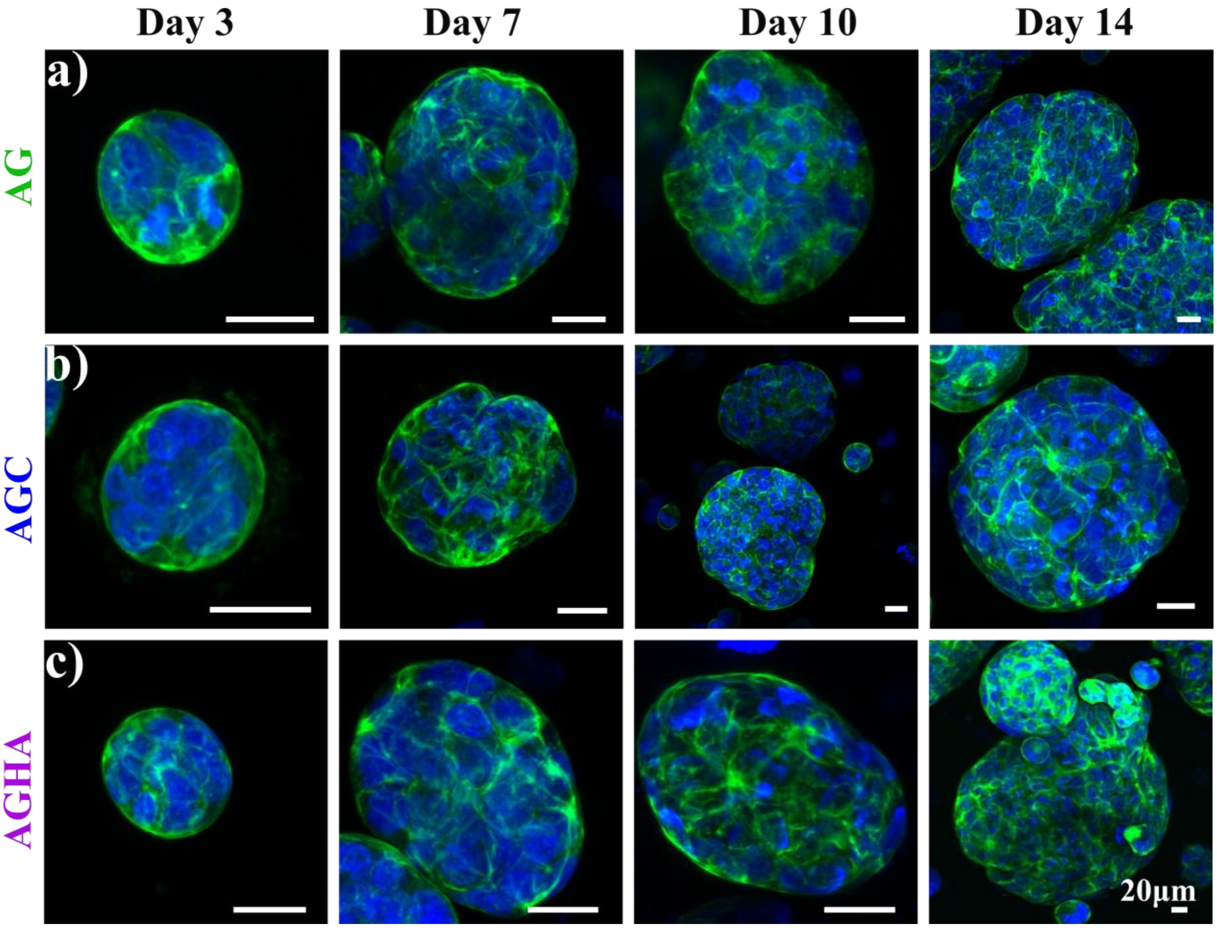


**Fig. 2**. Confocal imaging of F-actin distribution in NS-SV-AC spheroids growing in hydrogels. Cells were blended into a) AG, b) AGC, or c) AGHA gels and cultured for 14 days. Green, F-actin; blue, nuclear staining. Magnification ×20, scale bar 20 µm.

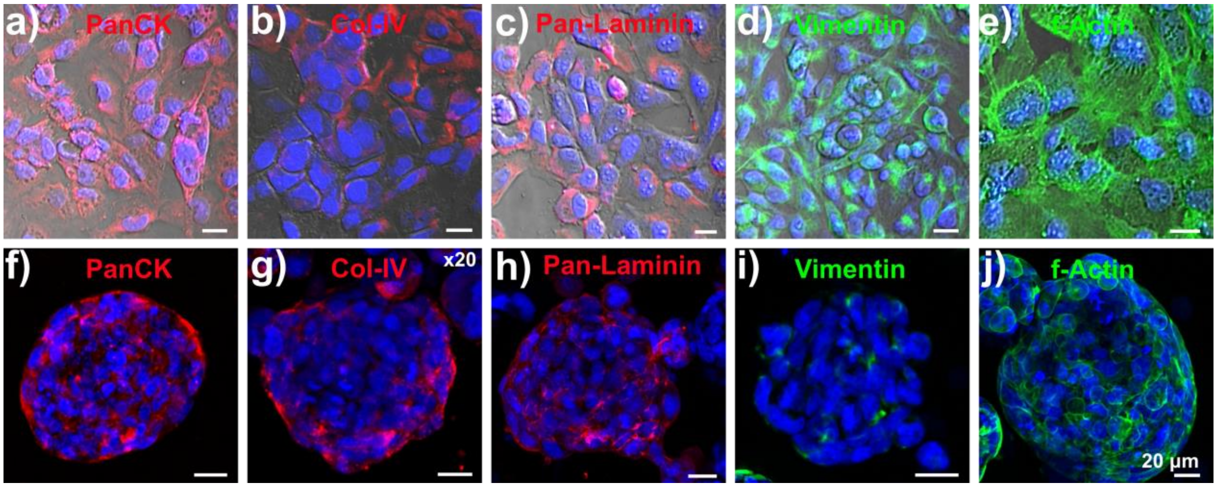


**Fig. 3**. Immunodetection of salivary gland markers in NS-SV-AC cells growing in 2D (4 days in culture) and in 3D platforms using AGHA 3D models (14 days in culture). 2D maximum projection of markers expressed by cells in 2D (a-e) or 3D (f-j) systems. For 2D culture images (upper panel), a bright field was included in the merged images to delimit individual cells. Red, PanCK, Col-IV, or Pan-Laminin; green, Vimentin or F-actin; blue, nuclear staining. Magnification x20, scale bar 20 µm.

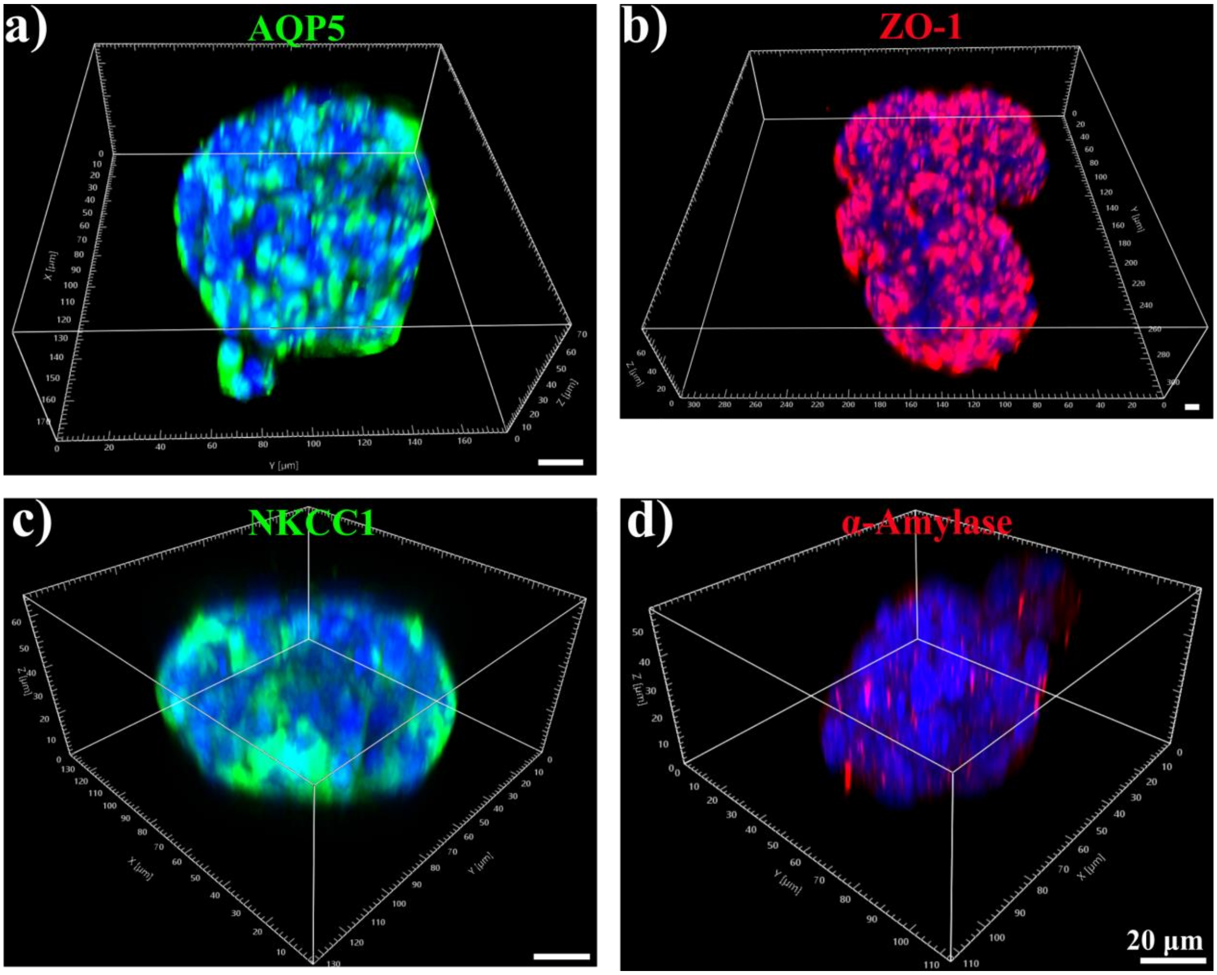


**Fig. 4.** 3D rendered images of NS-SV-AC spheroids positive to AQP5 (a), ZO-1 (b), NKCC-1 (c), and α-amylase (d) growing in AGHA hydrogels for 14 days. Green, AQP5 or NKCC-1; red, ZO-1 or α-amylase; blue, nuclear staining. Scale bar 20 μm.

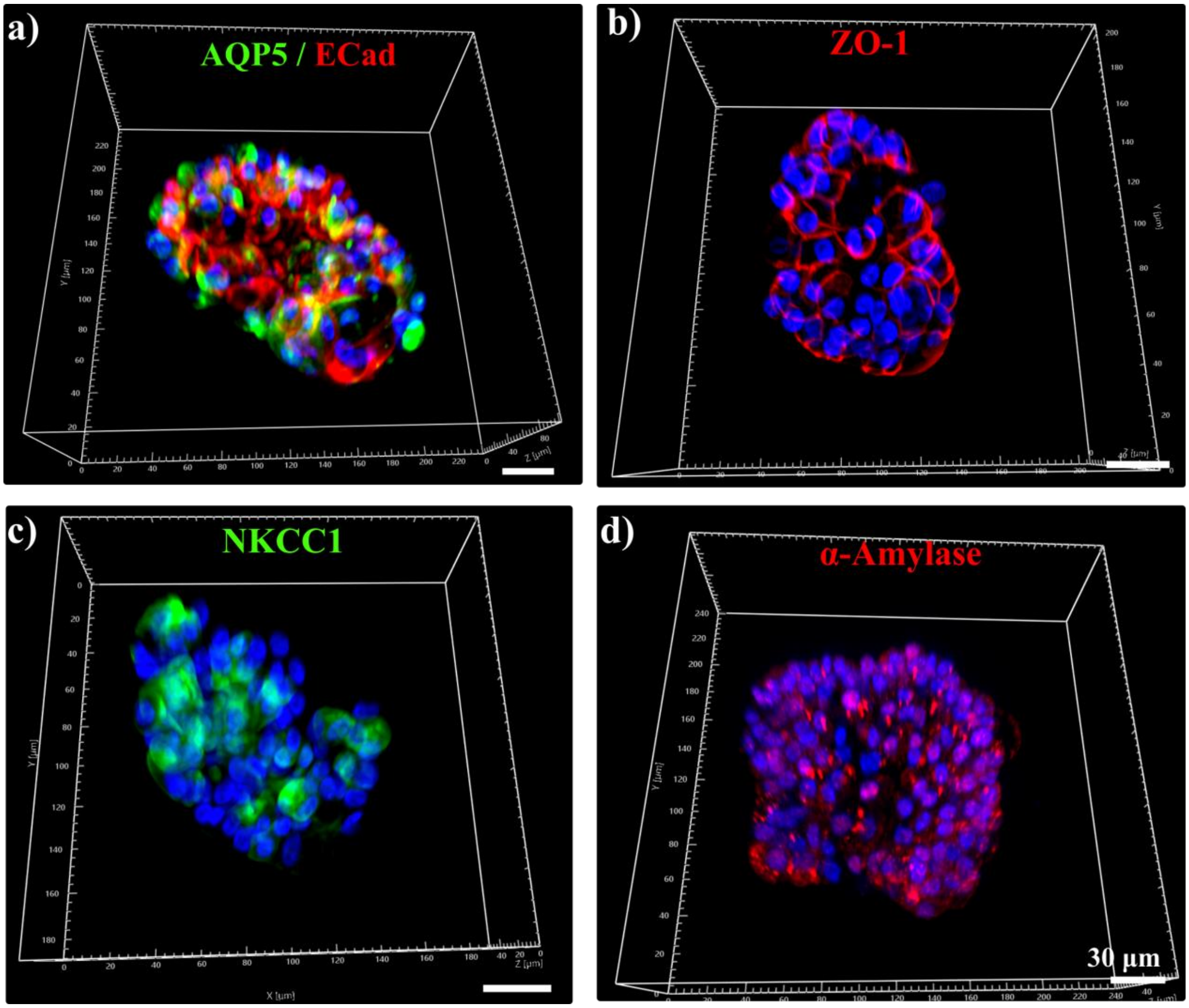


**Fig. 5**. 3D rendered images of SFUs positive to AQP5/ECad (a), ZO-1 (b), NKCC-1 (c), and α-Amylase (d) growing in AGHA hydrogel for 15 days. Green, AQP5 or NKCC-1; red, ECad, ZO-1, or α-amylase; blue, nuclear staining. Scale bar 30 μm.
